# *Pseudomonas aeruginosa* essential gene perturbations that confer vulnerability to the mammalian host environment

**DOI:** 10.1101/2024.11.05.622029

**Authors:** Neha K. Prasad, Ryan D. Ward, Michelle A. Yu, Michael S. Kwon, Amy B. Banta, Oren S. Rosenberg, Jason M. Peters

## Abstract

Multidrug-resistant *Pseudomonas aeruginosa* causes highly morbid infections that are challenging to treat. While antibiotics reduce bacterial populations during infection, the host environment also plays a key role in inhibiting and eliminating pathogens. Identifying genetic targets that create vulnerabilities to the host environment may uncover strategies to synergize with nutrient limitation or inherent immune processes to clear bacterial infections. Here, we screened a partial knockdown library targeting *P. aeruginosa* essential and conditionally essential genes in a murine pneumonia model to identify genes with increased vulnerability in the host environment. We found that partial CRISPR interference (CRISPRi) knockdown of 178 genes showed significant fitness defects in mice relative to axenic culture. We validated two important outliers: *ispD*, encoding a key enzyme in isoprenoid precursor biosynthesis, and *pgsA*, encoding an enzyme involved in phospholipid synthesis that is strongly upregulated in human infections. Partial knockdown of both genes showed decreased virulence in a mouse survival assay but had little impact on *in vitro* growth. The use of CRISPRi screening to uncover genetic vulnerabilities represents a promising strategy to prioritize antibacterial targets that interact with the host environment.

## Introduction

*Pseudomonas aeruginosa* is an environmental bacterium that is a common causative agent of both acute and chronic infections. Due to its inherent resistance to antibiotics and increasing levels of acquired resistance, multidrug-resistant *P. aeruginosa* has been prioritized as a high priority pathogen by the World Health Organization in 2024 (1). While *P. aeruginosa* is estimated to have 321 core essential genes required for growth of multiple strains under multiple culturing conditions (2), only a small fraction of these genes have been targeted for inhibition by small molecule antibiotics in clinical use and clinical development (3).

Although antibiotics are useful for reducing the bacterial burden during infection, their interactions and potential synergy with the native host environment for bacterial clearance are underexploited. It is likely that the extent of target inhibition required for bacterial growth inhibition *in vitro* may exceed that which is needed *in vivo* for specific targets, where the host environment including the immune system mediates clearance of the infection. For example, the synergy of beta-lactam antibiotics with host-produced antimicrobial peptides has been shown to reduce the burden of bacteria demonstrating *in vitro* resistance to the beta-lactam (4–8). Consequently, target-based whole-cell screens in antibiotic discovery efforts may neglect chemical matter with sufficient *in vivo* efficacy due to poor *in vitro* potency. Comparison of *in vitro* minimal inhibitory concentrations of antibiotics with their associated reduction of bacterial burden during *in vivo* infections is confounded by pharmacokinetic and pharmacodynamic properties of the antibiotic. Thus, a goal of bacterial geneticists has been to substitute chemical inhibition with genetic inhibition, thereby eliminating this confounding effect and expanding the scope of potential antibacterial gene targets to include those without known chemical inhibitors.

For genes that are non-essential *in vitro*, large-scale fitness assessment through transposon insertion sequencing has previously led to the classification of *in vivo* gene essentiality (9). Transposon sequencing of *P. aeruginosa* under various infection conditions has revealed many virulence factors, where gene knockout leads to attenuated virulence of the mutant strain (10). However, anti-virulence interventions have yet to demonstrate clinical efficacy and may not be suitable for people experiencing chronic *P. aeruginosa* lung infections associated with cystic fibrosis, which is often characterized by downregulation or loss-of-function mutations in virulence associated genes (11). Given that complete genetic inhibition strategies cannot be used to probe potential antibiotic targets that are essential *in vitro*, a partial genetic perturbation strategy enables us to probe this valuable category of genes.

Importantly, the notion of essentiality implies a binary effect of genetic inhibition on bacterial fitness, even though intermediary inhibition with chemical drugs indicates that the effect of target inhibition on fitness is, instead, a continuous variable. This gradient is captured by gene vulnerability (12), where partial genetic perturbation of essential genes can confer a quantifiable fitness defect that does not lead to a complete loss of viability. The significance and magnitude of gene vulnerability varies based on culture conditions and can be measured by depletion of the specific mutant from a pooled library (12–16). Essential genes with large *in vivo* vulnerabilities may represent a promising new class of antibacterial targets, since antibiotics must often be administered at high dosages that are capped by dose-limiting adverse effects, and corresponding inhibitors with no *in vitro* efficacy may have been previously overlooked.

Essential genes have been historically difficult to manipulate precisely, as they are requisite for pathogen survival. CRISPR interference (CRISPRi), where a catalytically inactive variant of the Cas9 nuclease (dCas9) sterically hinders RNA polymerase elongation leading to reduced transcription, is a powerful tool for loss of function screens. We have previously developed Mobile-CRISPRi, a modular and scalable platform to construct knockdown strains in a variety of pathogens (17–19), which enables us to detect gene vulnerability under *in vitro* and *in vivo* settings.

Previous work phenotyping essential and non-essential genes in mouse models of infection suffered from bottleneck effects or laborious library construction. An inducible CRISPRi screen of genes in the Gram-positive bacterium, *Streptococcus pneumoniae*, in a murine pneumonia model revealed new potential virulence factors and the non-essentiality of a potential antibiotic target that was considered essential *in vitro* (20). However, severe infection-associated bottlenecks limited screen robustness: while 31 genes were identified to have greater *in vivo* vulnerability, knockdown of only one non-essential gene (*purA*) was confirmed to attenuate virulence. Co-infection with influenza A virus overcame the infection bottleneck allowing for identification of additional virulence-associated genes. A more recent study used an inducible proteolytic degradation library of essential genes in *Mycobacterium tuberculosis* to study antibiotic-essential gene interactions in splenic and pulmonary murine infection model (21). Inducible degradation was achieved by systematically tagging the 3′ ends of essential genes with degradation tags (DAS) and expressing the Clp protease adaptor SspB by feeding mice doxycycline-containing chow. Although this strategy was effective at identifying drug-essential gene synergies, construction of such a library is extremely resource intensive, limiting adoption in other bacteria. Further, some essential genes are recalcitrant to 3′ tagging due to disruption of protein folding. In contrast, the technology and methodology developed in the present study provides a blueprint for overcoming infection bottlenecks that can be readily translated to many other pathogens and infection models.

Here, we developed a partial knockdown strategy that allowed us to probe the vulnerability of essential and conditionally essential *P. aeruginosa* genes *in vivo* (**Fig. 1**). We generated a pooled CRISPRi library of strains that showed little to no phenotype when grown in rich medium, then screened the library for *in vivo* vulnerability in a murine pneumonia model. Our approach enabled quantification and minimization of experimental bottlenecks by monitoring depletion of non-targeting control single guide RNAs (sgRNAs). We identified dozens of genes that are differentially vulnerable to the host environment, providing a resource for future therapeutic efforts. Finally, we follow up two intriguing hits, *ispD* and *pgsA*, showing that perturbation of these genes outside the pooled context allows for survival of mice that would otherwise succumb to infection with wild-type *P. aeruginosa* at the same dose. The simplicity of our strategy suggests that it will be broadly applicable.

**Figure 1.**
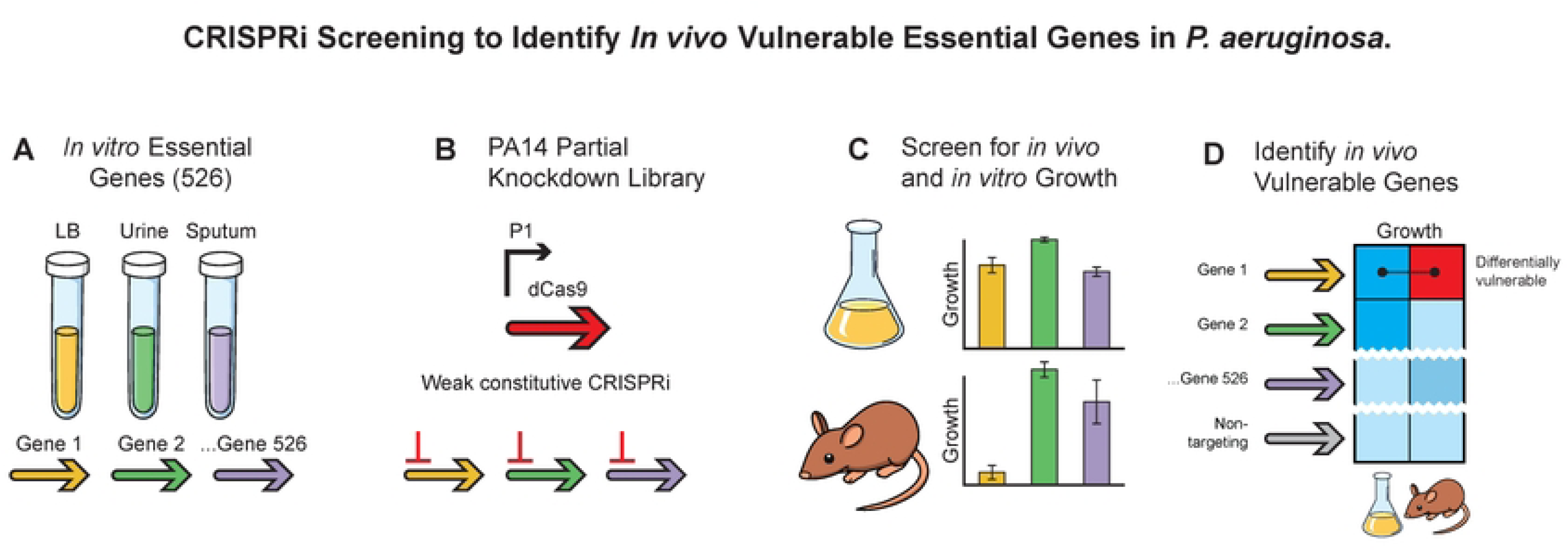
Overview of the CRISPRi screening approach for identifying in vivo essential genes in *P. aeruginosa* PA14. **(A)** Essential genes are identified from literature sources based on their necessity for growth in various media, including LB, urine, and sputum, resulting in 526 essential genes. **(B)** A partial knockdown library for PA14 is constructed using Mobile-CRISPRi with a weak constitutive promoter (P1) to partially inhibit these essential and conditionally essential genes. **(C)** The library is screened for growth effects both in vitro and in a murine pneumonia model, allowing comparison of gene essentiality across different environments. **(D)** Genes that are differentially vulnerable in the host versus LB growth conditions are identified, depicted as a heatmap to illustrate variations in gene vulnerability.

## Results

### Pooled Construction of a *P. aeruginosa* Essential Gene Knockdown Library

To mitigate concerns pertaining to the extent and uniformity of gene knockdown when using an inducible CRISPRi promoter in an *in vivo* model, we previously characterized the efficacy of constitutive promoters expressing dCas9 to drive CRISPRi knockdown (17–19). We elected to use weakest promoter (P1 resulting in ∼10-fold knockdown) for our essential gene knockdown library with the goal of causing a modest perturbation that would show limited fitness effects *in vitro* but potentially large effects *in vivo*.

We selected 526 genes to target in our *P. aeruginosa* PA14 essential gene knockdown library based on a transposon sequencing study that identified the “core” essential genome for *Pseudomonas*. (2) Genes that were deemed essential in at least one of the lab- or infection-related growth media (lysogeny broth (LB), minimal medium (M9), synthetic Cystic Fibrosis sputum (SCFM), fetal bovine serum (FBS), and urine) were included in our library design. For each gene in the library, we designed four non-overlapping sgRNA spacers targeting the 5′ end of the coding sequence (**Supplementary Table S1**), (18). To assess bottlenecks and control for the effects of CRISPRi knockdown during *in vivo* experiments, we included 1,000 non-targeting sgRNAs in the library as negative controls. Following cloning of the pooled spacer library, CRISPRi vectors were transferred to WT PA14 via triparental mating and were chromosomally integrated into *att*_Tn*7*_ site (**Supplementary Figure 1**), (17–19).

We quantified the representation of non-targeting sgRNAs using Illumina next generation sequencing (NGS), finding a normal distribution of spacer counts in the pooled mating strain library and in the pooled PA14 knockdown library (**Supplementary Fig. 2A**). This suggests that there were no substantial technical bottlenecks in construction of the mating strain library for fitness-neutral guides. However, NGS revealed that 10 genes were not detected in the knockdown library after transformation into PA14 (**Supplementary Fig. 2A**), possibly due to excessive knockdown for highly sensitive genes. Ultimately, the PA14 knockdown library targeted 516 genes represented by at least one sgRNA (**Supplementary Fig. 2B**).

### An *in vivo* CRISPRi screen in *P. aeruginosa* murine pneumonia model overcomes infection-associated bottlenecks

Pooled library infections are affected by bottlenecks that can confound the effects of genetic inhibition on measurement of strain loss after infection (20,22). Bottlenecks can arise from several mechanisms, such as physical barriers to infection, strain loss during inoculation, host clearance pathways as the bacteria transition to invasive disease, or stochastic depletion of mutant strains. Other technical issues with experimental infection models in animals include induction of fatal septic shock with too high of a bacterial inoculum, underrepresentation of the library at time of inoculation or sacrifice, and insufficient duration of infection resulting in too few bacterial doublings–all of which disallow robust detection of strain depletion.

We sought to address these pitfalls in our experimental approach (**Fig. 2A-C**). First, we used direct intratracheal instillation (23), as we reasoned that it would outperform indirect delivery methods (*e.g.,* intranasal instillation) which may generate an additional physical barrier and subsequent strain loss. Second, we minimized the loss of knockdown strains targeting highly vulnerable genes by inoculating mice with a dilution from a thawed glycerol stock, rather than allowing the pooled libraries to grow in axenic culture before inoculation. Third, we amplified sgRNA spacers for library quantification from *P. aeruginosa* colonies that grew after plating lung homogenates to avoid PCR issues that arose from attempting to amplify directly from the lung homogenates (**Fig. 2B**). Finally, as a strategy to distinguish unique vulnerabilities associated with the infection environment from general growth defects conferred by repression of an essential gene, we carried out an *in vitro* screen in parallel to the *in vivo* screen (**Fig. 2C**).

**Figure 2.**
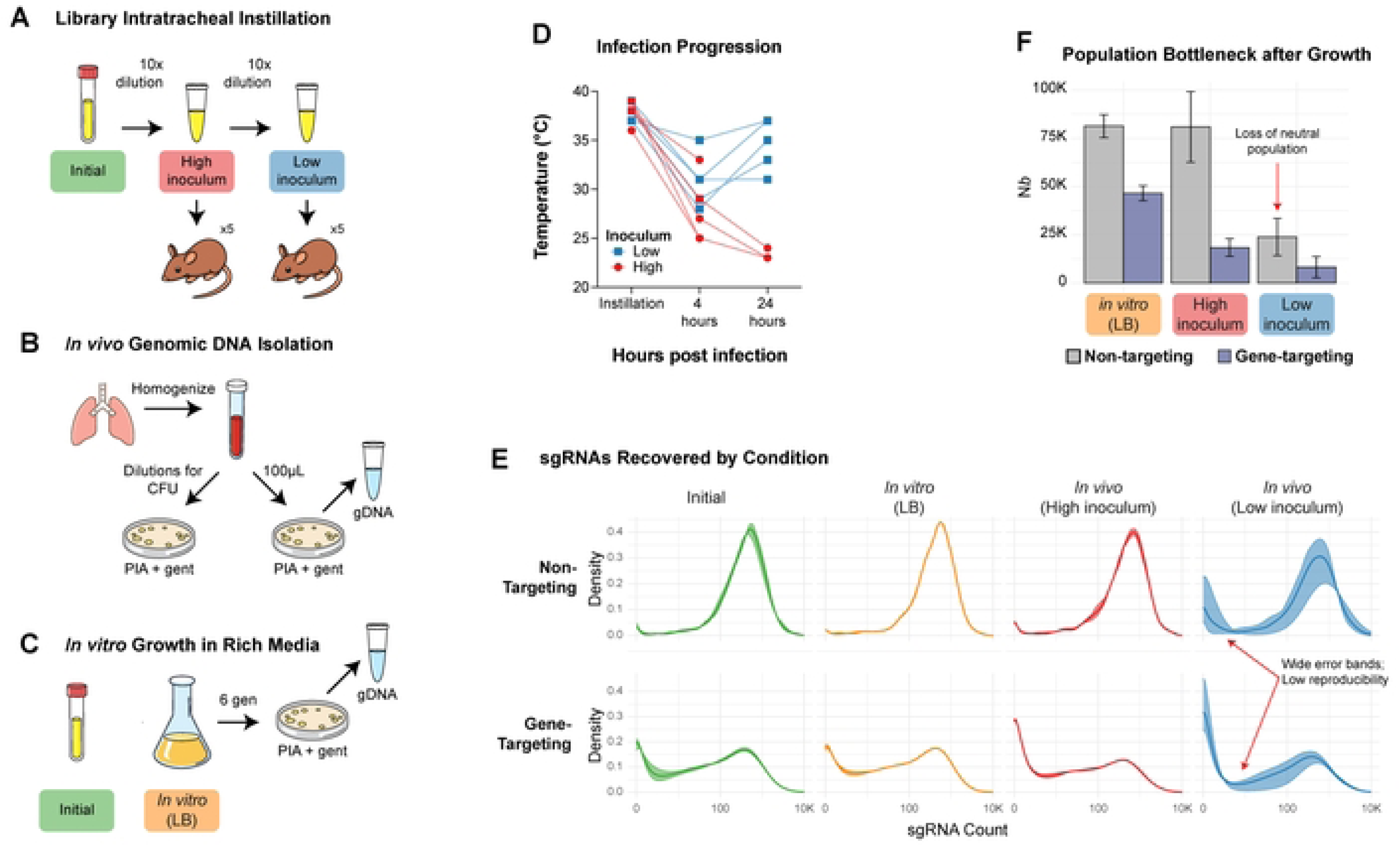
Murine Pneumonia Infection with the PA14 Essential Gene Knockdown Library. **(A)** Two groups of mice (n=5 per group) were intratracheally instilled with the PA14 knockdown library at high (4.6 × 10¹¹ CFU/animal) or low (4.6 × 10¹⁰ CFU/animal) doses to assess infection dynamics. **(B)** After 24 hours of infection, lung homogenates were plated on *Pseudomonas* isolation agar (PIA) containing 30 μg/mL gentamicin. Colonies were harvested after 24 hours, and genomic DNA was extracted for amplicon library preparation and sequencing. **(C)** Concurrently, inoculum samples were cultured in 25 mL of LB medium and grown for six generations before plating on PIA with gentamicin, allowing comparison between in vivo and in vitro growth dynamics. **(D)** Individual changes in mouse rectal temperatures and body weights over 24 hours post-infection to assess physiological impacts. Each line represents data from a single animal. **(E)** Distribution of single guide RNA (sgRNA) counts recovered from the inoculum, *in vitro* culture, and lung homogenates from high and low inoculum cohorts, illustrating sgRNA representation across experimental conditions. **(F)** Population bottleneck analysis comparing non-targeting and gene-targeting constructs across *in vitro* growth and both high and low inoculum infection conditions. All plotted data show mean ± SEM, except for panel D, which represents individual animals.

To ensure an appropriate *P. aeruginosa* inoculum for our acute infection experiments, two groups of five mice were intratracheally instilled with approximately 4.6E11 CFU/animal (high inoculum) and 4.6E10 CFU/animal (low inoculum) of the PA14 knockdown library (**Fig. 2A**). This 1-log variation in the bacterial inoculum drastically affected the ability of mice to clear the infection, yielding a screening bottleneck for the low inoculum group. In the low inoculum group, the recovery of animal temperature and weights by the 24-hour time point indicated drastic clearance of infection (**Fig. 2D**). Indeed, the population of negative control sgRNAs was skewed in the lung homogenate samples recovered from this group, suggesting stochastic depletion independent of genetic perturbation (**Fig. 2E-F**). NGS accordingly exposed bottlenecks in the samples recovered from mice infected with the diluted inoculum that prohibited downstream assessment of gene vulnerability.

In contrast, this screening bottleneck was not apparent within the group of mice infected with the more concentrated inoculum. Two of the five mice succumbed to the infection during the 24-hour period, which may be attributable to septic shock, based on the relatively low weight loss coupled with large temperature change in the three surviving mice (**Fig. 2D**). In the lung homogenates from the three surviving mice, approximately log-normal distributions of non-targeting controls were recovered, implying the absence of a substantial strain bottleneck (**Fig. 2E**). We evaluated the bottleneck size by calculating the population complexity of the library screened under axenic or *in vivo* conditions, finding that complexity of the non-targeting controls was similar under both conditions. (**Fig. 2F**). The similarity of the distributions between the inoculum and *in vitro* samples suggests that the least fit strains in the PA14 knockdown library had already been depleted during the library construction process (**Fig. 2E**). In contrast, the distribution of gene-targeting sgRNAs is more skewed in the *in vivo* sample, with a larger fraction of sgRNAs showing fewer counts, indicating that the fitness of many knockdown strains is reduced in the infection environment (**Fig. 2E**).

### An *in vivo* CRISPRi Screen Reveals Gene Vulnerability during Murine Pneumonia

In probing the importance of PA14 genes during murine pneumonia infection, we sought to identify hypomorphs with heightened vulnerability to clearance by the host. As the vast majority of genes in the library were targeted by multiple sgRNAs, we calculated the median extent of depletion of the corresponding knockdown strains to determine which essential gene perturbations conferred vulnerability to the mammalian host infection environment relative to nutrient-rich growth medium. Of the 516 genes targeted in the PA14 knockdown library, we were able to calculate changes in relative abundance for 466. Strains corresponding to 178 genes were depleted (Log_2_ Fold Change (LFC) < -1, False Discovery Rate (FDR) < 0.05) after 24 hours of growth in the mouse lung compared to cells grown in axenic culture (**Fig. 3A,B**). In contrast, no significant *in vitro* vulnerabilities were detected when compared to the distribution of PA14 knockdown library strains in the inoculum, consistent with the idea that knockdown strains with strong negative phenotypes were largely absent from the starting PA14 library. (**Fig. 3B**). The lack of detectable genetic vulnerabilities during *in vitro* growth supports the notion that strains in the library inoculum are fit for growth in a nutrition-rich environment.

**Figure 3.**
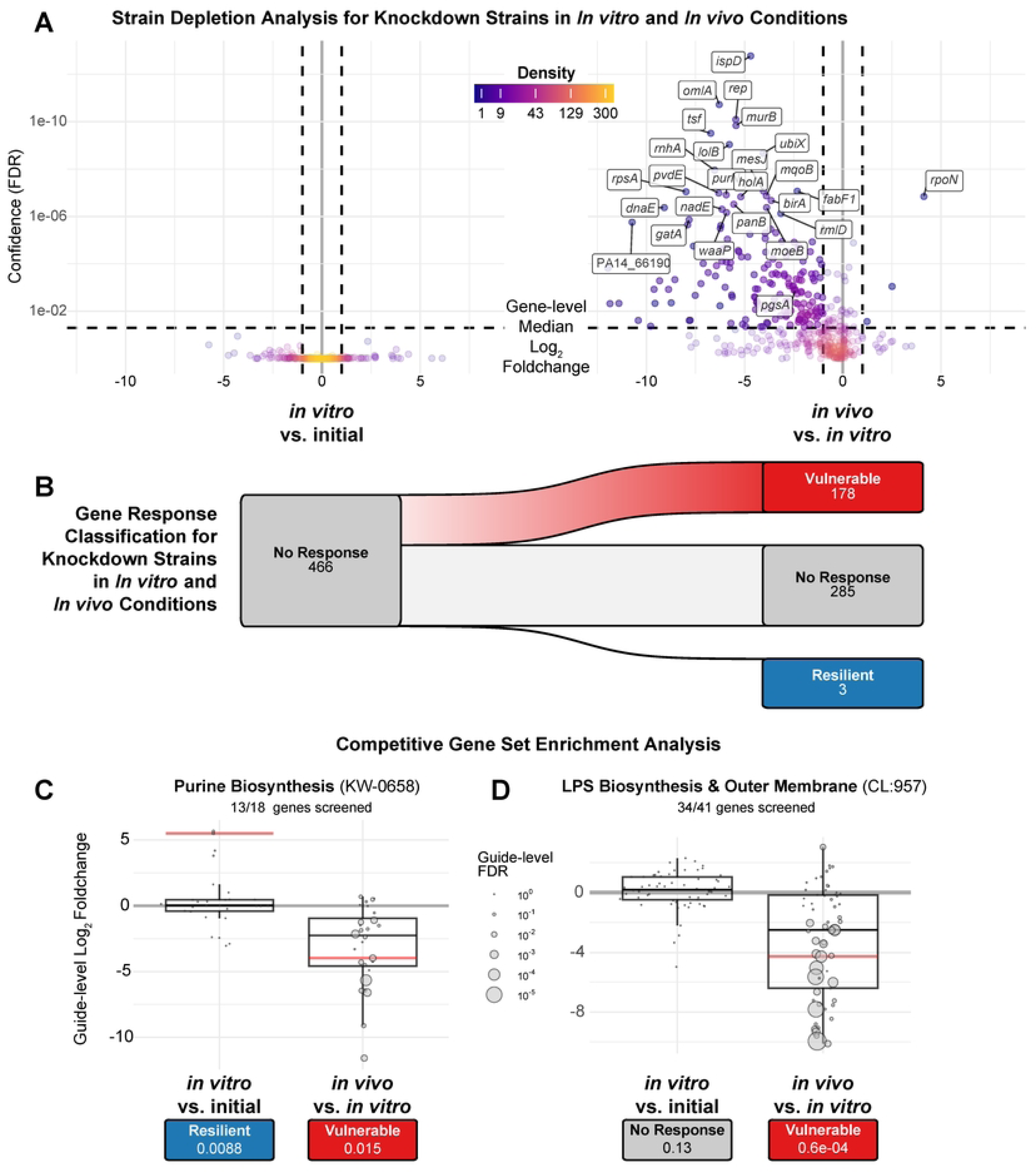
PA14 Essential Gene Vulnerabilities Identified in In Vitro and In Vivo Screens. **(A)** Volcano plots representing changes in strain composition for knockdown strains across two key comparisons: *in vitro* vs. initial (growth effects) and *in vivo* vs. *in vitro* (additional effects in mouse lung environment). Color intensity indicates sgRNA density, with darker colors representing higher density. The y-axis shows FDR (false discovery rate) on an inverted scale, and the x-axis shows the median log_2_ fold change for each targeted gene. **(B)** Gene response classification for knockdown strains in *in vitro* and *in vivo* conditions. The Sankey diagram illustrates transitions between response categories: “Vulnerable” (red, FDR ≤ 0.05 and median log_2_ fold change < -1), “Resilient” (blue, FDR ≤ 0.05 and median log_2_ fold change > 1), and “No Response” (gray, FDR > 0.05). Width of connecting bands indicates the number of genes in each transition. (**C**, **D**) Competitive gene-set enrichment analysis using CAMERA for (**C**) Purine Biosynthesis and (**D**) LPS (Lipopolysaccharide) Biosynthesis & Outer Membrane pathways. Log2 fold change for guides targeting genes in these pathways is shown for *in vitro* vs. initial and *in vivo* vs. *in vitro* comparisons, with FDR values noted below each condition. Boxplots show median (center line), interquartile range (box), and 1.5× interquartile range (whiskers). Individual points represent individual guides, with point size reflecting guide-level significance (FDR). Red lines indicate the significance-weighted median for significantly enriched gene sets (FDR ≤ 0.05).

We next investigated *in vivo* vulnerabilities at the pathway level using both gene-level and guide-level analyses. Our initial attempts to aggregate sgRNAs at the gene level failed to identify biological pathways that were enriched for vulnerability in vivo, likely due to increased noise of CRISPRi screening in the host environment relative to axenic culture. To overcome this limitation and better distinguish true signals from noise, we adopted a more nuanced approach. We analyzed all sgRNAs targeting genes within each pathway collectively, effectively increasing the number of measurements we could use to calculate pathway-level statistics. Using this guide-level analysis and competitive gene-set enrichment analysis (CAMERA), we identified several biological pathways that were significantly perturbed and well-represented in the knockdown library—specifically, pathways where at least 70% of the genes were targeted by sgRNAs (**Supplementary Table S3**). Notably, pathways involved in folate biosynthesis (KW-0289) and isoprene biosynthesis (CL:924) exhibited significant *in vivo* vulnerabilities when compared to the *in vitro* baseline, whereas arginine biosynthesis (CL:2388) exhibited *in vivo* resilience (**Supplementary Fig. 3**). Additionally, the purine biosynthesis pathway (KW-0658) and the lipopolysaccharide biosynthesis and outer membrane assembly pathway (CL:957) also showed significant depletion *in vivo* (**Fig. 3C,D**). This guide-level approach allowed us to detect critical pathways that may not have been identified through traditional gene-level analyses.

As expected, purine biosynthesis showed significant vulnerability at the pathway level (**Fig. 3C**). Previous transposon insertion sequencing (Tn-seq) studies showed that multiple purine biosynthesis genes (*purD*, *purE*, *purF*, *purH*, *purK, purL*, *purN*) are dispensable for growth in rich medium but are required in synthetic sputum (SCFM), whereas *purA* and *purB* are essential in both media (2). A previous study also revealed the essentiality of *purA* in a *S. pneumoniae* murine pneumonia model (20). In our CRISPRi screen, significant gene-level vulnerability was detected for *purA*, *purD*, *purF*, *purL*, *purN*, but not for *purB*, *purK*, *purL*, and *purN*. Given that transposon disruption of *purD*, *purF*, *purL*, and *purN* does not significantly impede bacterial growth in rich media, inhibitors that exploit this *in vivo* vulnerability may not exhibit antibacterial activity in conventional assays, complicating development efforts.

Similarly, CRISPRi knockdown of lipopolysaccharide (LPS) biosynthesis and outer membrane related genes showed significant *in vivo* vulnerability at the pathway level (**Fig. 3D**). Tn-seq studies have shown that all seven genes of the LPS transport complex (*lptA*, *lptB, lptC, lptD, lptE, lptF,* and *lptG*) are essential in both rich media and SCFM (2). Specifically, *lptC*, *lptD*, *lptF*, and *lptG* demonstrated significant vulnerability in the mouse lung environment, while *lptA*, *lptB*, and *lptE* did not show this context-specific depletion. In agreement with these findings, a conditional deletion of *lptA* (a.k.a., *lptH*) in PA01 has previously been shown to attenuate virulence in a murine pneumonia model.^24^ Partial genetic perturbation allows us to build upon such observations and identify other genes with heightened *in vivo* vulnerability.

The gene encoding the alternative σ factor, RpoN (*rpoN*), is a rare example of a knockdown that decreases *in vivo* vulnerability (**Fig. 3B**). RpoN has been shown to regulate virulence pathways (24) and deletion of *rpoN* results in 100-fold less virulence in a mouse thermal injury model (25). However, *P. aeruginosa* commonly evolves *rpoN* loss-of-function mutations during chronic infection of cystic fibrosis patients, suggesting that loss of RpoN activity may also improve strain fitness in certain contexts (11,26,27). Modifications of pathogen-associated molecular patterns (PAMPs), such as those linked to RpoN regulation, enable immune evasion and survival in the infection environment through hindering immune recognition and activation (28,29). We speculate that virulence-related activities reduced by *rpoN* knockdown are effectively complemented by other strains in the pooled environment, allowing RpoN-deficient cells to escape host clearance mechanisms.

### Validation of Knockdown Vulnerability in Murine Pneumonia Model

Next, we sought to validate hit genes from the pooled screen through assessing survival rates of mice infected with individually constructed knockdowns. Validation with individual knockdown strains could reveal any dependencies of the hypomorphic strain’s *in vivo* clearance on the presence of other constituents in the PA14 essential gene knockdown library. For example, deficiencies in one member of the library may only be partially compensated for by other co-infected members of the library in the case of virulence pathways mediated by secreted products, like siderophore production, or community dynamics (30). We pursued two independent strategies to select hypomorphic strains for validation studies: 1) strains that exhibited significantly greater *in vivo* vulnerability than *in vitro* vulnerability; 2) strains corresponding to core essential genes that were previously found to be upregulated during human infection.

Under the first strategy, *ispD* stood out as the most confident hit and was selected for validation studies (**Fig. 3B**). As part of the methylerythritol phosphate (MEP) pathway towards the non-mevalonate biosynthesis of isoprenoid precursors, IspD (2-C-methyl-D-erythritol 4-phosphate cytidylyltransferase) catalyzes the formation of 4-diphosphocytidyl-2-C-methyl-D-erythritol. Given most Gram-negative bacterial species use this non-mevalonate pathway whereas humans exclusively use the mevalonate pathway, IspD represents a potential antibacterial target that is currently unexploited in the clinic (31). While fosmidomycin is a known inhibitor of the MEP pathway through inhibition of IspC (32), few IspD inhibitors with antimicrobial activity have been reported– limited to *Plasmodium falciparum* (33,34), *Acinetobacter baumannii* (35), and biodefense pathogens *Yersinia pestis* and *Francisella tularensis* (36).

Under the second strategy, *pgsA* stood out as the most confident hit and was selected for validation studies. As part of the selection process, we compared our set of *in vivo* vulnerable genes to a previously published dataset of genes known to be strongly upregulated during human infection (37). Previous studies revealed little overall correlation across the entire genome between transcriptionally important genes, whose expression is affected by a change in the environment, and phenotypically important genes, whose fitness is affected by a change in the environment (38). However, phenotypically and transcriptionally important genes were found to overlap when probing the effects of nutritional stress on metabolic genes (38). We hypothesized that core essential genes that are upregulated during infection will be vulnerable in the host if they cannot be upregulated due to genetic or chemical inhibition, and thus may represent promising antibiotic targets. Based on published datasets, we found that only 4 genes were both “core” essential (2) and significantly upregulated (>2 LFC) during human infection (37): *lptA, lptG, pgsA, cysS*. As previously mentioned, *lptA* and *lptG* are part of the lipopolysaccharide transport system, and a conditional deletion of *lptA* in *P. aeruginosa* has previously been shown to have attenuated virulence in a murine pneumonia model (39). The remaining two genes, *pgsA* and *cysS*, are involved in phospholipid biosynthesis and tRNA aminoacylation, respectively. Of the four genes, only *lptG* and *pgsA* exhibited significant vulnerability when comparing *in vivo* to *in vitro* conditions, indicating specific depletion in response to the mouse lung environment (**Supplementary Table S2**). Since *pgsA* was observed to be the most significantly upregulated of the four during human infections, the corresponding knockdown was chosen for validation studies.

Genetic knockdown mutants of *ispD* and *pgsA*, as well as a control non-targeting strain (NTC), were generated with Mobile-CRISPRi using the same P1 constitutive promoter from the pooled library. As expected, growth of the *ispD* and *pgsA* partial knockdown mutants in LB media was not substantially different from either WT PA14 or the non-targeting control strain (**Fig. 4A**), demonstrating the lack of *in vitro* fitness defects.

**Figure 4.**
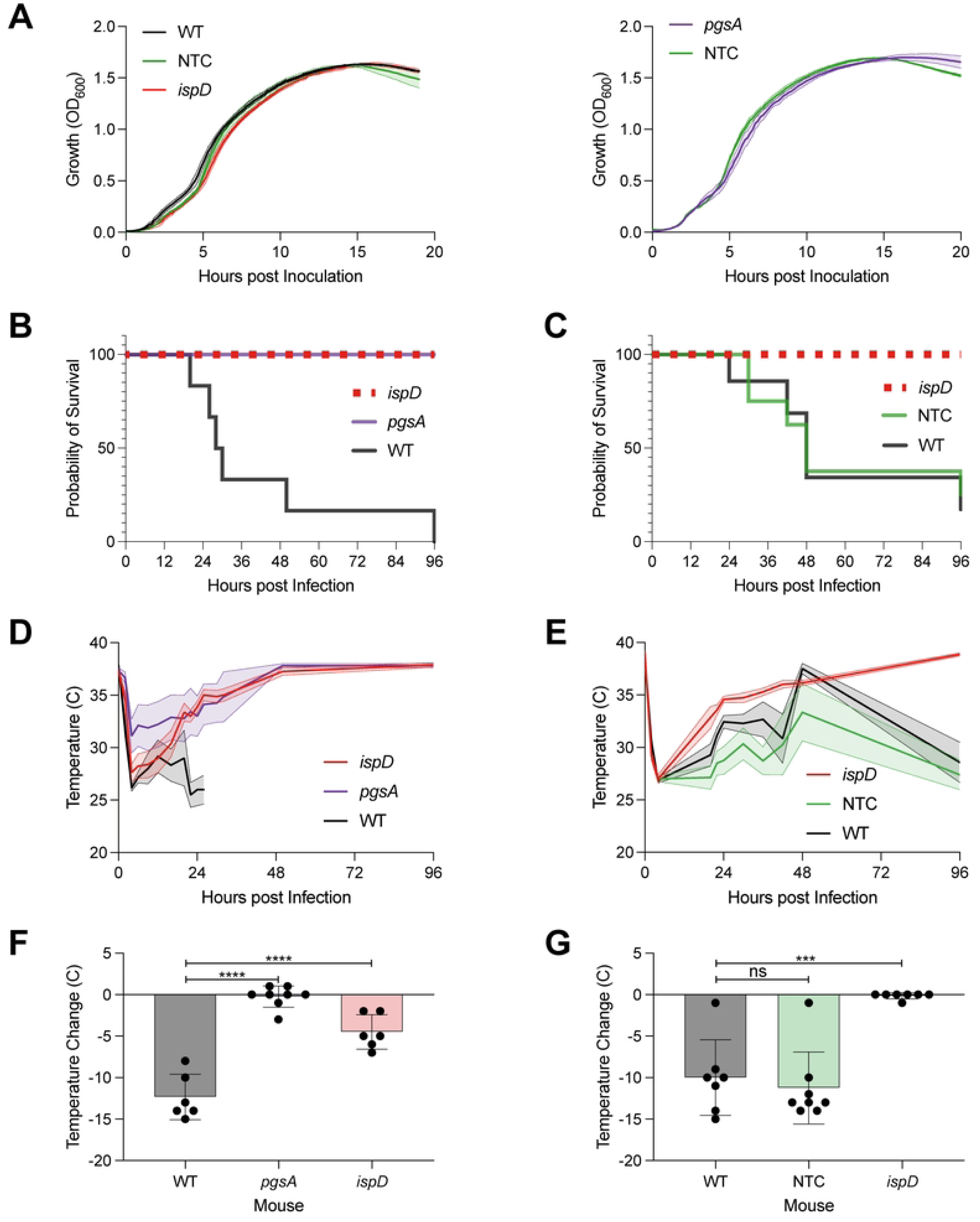
Effects of *ispD* and *pgsA* Knockdowns on Virulence and Mouse Thermoregulation. **(A)** In vitro growth curves of wild-type PA14, non-targeting control (NTC), *ispD* knockdown mutant, and *pgsA* knockdown mutant. Lines represent the mean, and shaded areas represent SEM across biological replicates. **(B, C)** Survival curves of mice infected with wild-type PA14, *pgsA* knockdown, *ispD* knockdown, or NTC mutants. **(D, E)** Changes in body temperature of mice infected with the indicated strains. Lines represent mean values, and shaded areas indicate SEM. **(F, G)** Temperature loss at 20 hours post-infection with the indicated strains. Data are presented as mean ± SEM for 6 to 8 mice per group. Statistical analysis was performed using one-way ANOVA compared to wild type. \*\**p* ≤ 0.001; \*\*\**p* ≤ 0.0001.

To validate our screen and confirm that *ispD* and *pgsA* knockdowns were defective for pathogenesis outside of the pooled context, we monitored the survival of mice infected with *ispD* and *pgsA* hypomorphs individually. We intratracheally instilled planktonic monocultures of the *pgsA* and *ispD* knockdown mutants as well as the negative control strain and WT PA14 at 2E6 CFU/animal. Mice infected with the *ispD* and *pgsA* mutants showed 100% survival over 4 days, while the mean survival time of mice infected with WT PA14 or the non-targeting control strain was substantially lower (**Fig. 4B and 4C**). This difference in pathogenicity is also reflected by the observation that mice instilled with *ispD* or *pgsA* knockdown strains more rapidly recovered their body temperature post-infection (**Fig. 4D and 4E**), including a statistically significant difference in temperature loss at the 20-hour time point (**Fig. 4F and 4G**). These findings suggest that partial inhibition of the core essential genes *pgsA* and *ispD* significantly enhances infection clearance *in vivo,* despite the lack of noticeable phenotypic effects *in vitro*.

## Discussion

Our pooled *in vivo* CRISPRi screen reveals a heightened vulnerability to the host environment for 178 PA14 genes, providing a genetic resource for promising therapeutic targets. In particular, genetic perturbation of the essential genes *pgsA* and *ispD* did not significantly affect the growth of PA14 in axenic culture, but the same perturbation *in vivo* impeded productive host infection by these mutants. Given that these bacterial genes are required for growth in a mammalian host and are not conserved in humans, they may have high therapeutic potential.

Perturbation of both *pgsA* and *ispD* would both be expected to disrupt envelope functions in *P. aeruginosa*: *pgsA* by affecting phospholipid composition (40) and *ispD* by affecting the levels of isoprenoid precursors available for peptidoglycan synthesis (41). However, the precise mechanisms of how this disruption would cause increased sensitivity *in vivo* are unknown. Connections between bacterial isoprenoids and the human immune system are well established, suggesting that immune detection could play a role (42). Vγ9/Vδ2 T cells are known to detect (E)-4-Hydroxy-3-methyl-but-2-enyl pyrophosphate (HMB-PP), a MEP pathway intermediate produced by the enzyme IspG (43). This special subset of T cells rapidly responds to the presence of microbes in human blood (42). As IspD lies upstream of IspG in MEP synthesis, the expectation would be that *ispD* knockdown would ultimately produce less HMB-PP and, therefore, reduce stimulation of Vγ9/Vδ2 T cells. Perhaps the potential effects on membrane disruption outweigh any potential immune evasion gained from reducing HMB-PP production. Future work will investigate the impacts of *pgsA* and *ispD* on cell physiology and membrane integrity.

There are limitations to the experimental design of this study. Firstly, the vulnerability profiles are dependent on the extent of knockdown elicited by the P1 promoter and consistent efficacy across the sgRNAs targeting each gene. Stronger inhibition may reveal vulnerabilities for other genes that were not significantly sensitive to repression driven by P1 in either *in vitro* or *in vivo* conditions. Conversely, weaker inhibition may alleviate growth-hampering fitness defects that may have led to underrepresentation of essential gene knockdown mutants in the inoculum. Secondly, comparison of *in vitro* fitness defects to *in vivo* fitness defects is inherently affected by the number of doublings that proceed under both conditions. Since the depletion of an unfit strain is expected to increase in magnitude over time, the fitness defect may not be detected if too few doubling times are captured. The *in vitro* 6-generation time point (OD_600nm_ 0.01 to 0.64) was chosen to mimic antibacterial discovery platforms, which typically track bacterial growth from a log-phase culture diluted to 1E5–1E6 CFU/mL until stationary phase. However, *in vivo* generation time likely does not match *in vitro* generation time, further complicating the comparison. Lastly, loss of function hypomorphs generated by gene knockdowns do not precisely mimic the effects of small molecule antibiotics, which may produce effects more similar to specific genetic alleles (e.g., loss of function in a specific domain or gain of function).

Importantly, the genetic vulnerability insights gleaned in this study may be a proxy for *P. aeruginosa* vulnerability to chemical inhibitors, revealing possible targets for small molecule inhibitors. Partial perturbation of genes corresponding to enzymes with natural substrates is akin to non-competitive chemical inhibition, as target depletion is equivalent to reducing V_max_ while leaving K_m_ unchanged (44). Furthermore, emerging therapeutic modalities such as CRISPR systems, targeted protein degradation, or antisense technology closely mimic the results of our partial genetic inhibition approach. However, just as the development of non-traditional antibacterials targeting virulence pathways has proven challenging due to the lack of *in vitro* MICs, inhibitors of genes with enhanced *in vivo* vulnerability may face similar barriers to drug development. Developing *in vitro* assays or identifying non-mammalian model organisms that are predictive of exploiting *in vivo* vulnerabilities (45) is critical for capitalizing on this paradigm of target prioritization.

The phenomenon where greater vulnerability is observed *in vivo* than *in vitro* for certain genes suggests that *in vitro* growth inhibitory measurements may undervalue the therapeutic potential of inhibiting these genes *in vivo*. Considering that many small molecule antibiotics have dose-limiting toxicities that have stymied their clinical development (3), the concept of achieving high efficacy of bacterial clearance with a reduced drug dose is especially pertinent.

## Acknowledgements

We thank the Chan-Zuckerberg Biohub for assistance with Illumina sequencing.

## Author Contributions

Conceptualization; Neha K. Prasad, Michelle A. Yu, Oren S. Rosenberg, Jason M. Peters

Methodology; Neha K. Prasad, Ryan D. Ward, Michelle A. Yu, Michael S. Kwon, Amy B. Banta, Oren S. Rosenberg, Jason M. Peters

Formal analysis; Neha K. Prasad, Ryan D. Ward, Michelle A. Yu, Michael S. Kwon, Jason M. Peters

Writing; Neha K. Prasad, Ryan D. Ward, Oren S. Rosenberg, Jason M. Peters

Funding acquisition; Oren S. Rosenberg, Jason M. Peters

Supervision; Michelle A. Yu, Oren S. Rosenberg, Jason M. Peters

## Resource Availability

### Lead Contact

Requests for further information and resources should be directed to and will be fulfilled by the lead contact, Jason Peters.

### Materials availability

The generated in this study will be made available on request, but we may require a payment and/or a completed materials transfer agreement if there is potential for commercial application.

### Data and code availability

The DNA sequencing data have been deposited at NCBI BioProject Database as PRJNA1178310 and will be publicly available as of the date of publication. During review, these data can be accessed using the following reviewer link: https://dataview.ncbi.nlm.nih.gov/object/PRJNA1178310?reviewer=q9ddvis29bnag1jcldcj6u7mng

All original code is openly available at GitHub (https://github.com/ryandward/pseudomonas_analytics/).

Any additional information required to reanalyze the data reported in this paper is available from the lead contact upon request.

## Materials and Methods

### Strains and Growth Conditions

*Escherichia coli* and *Pseudomonas aeruginosa* were grown in lysogeny broth (LB), Lennox (BD240230; 10 g tryptone, 5 g yeast extract, 5 g NaCl per liter) at 37°C aerobically in a flask with shaking at 250 rpm or in a culture tube on a roller drum. For growth on plates, LB was solidified with 1.5-2% agar (BD 214530). Antibiotics were added when necessary: *E. coli* (100 µg/ml ampicillin or carbenicillin, 15 µg/ml gentamicin) and *P. aeruginosa* (30 µg/ml gentamicin). Diaminopimelic acid (DAP) was added at 300 µM to support growth of DAP-auxotrophic *E. coli* strains. All strains were preserved in 15% glycerol at -80°C. Strains are listed in Supplementary Table 4.

### Mobile-CRISPRi Individual Gene and Gene Library Construction

sgRNAs were designed to knockdown essential genes (4 guides/gene plus 1000 non-targeting controls, 3112 total) in *P. aeruginosa* PA14 using a custom python script and *P. aeruginosa* genome sequences; RefSeq accession numbers NC_002516.2 and NC_008463.1 (18). Gene knockdowns included in the library were selected based on essential and conditionally essential genes defined in multiple growth conditions including LB, M9 minimal media, sputum, serum, and urine, with targeted genes listed in Supplementary Table S4 (2). sgRNA-encoding sequences were cloned into Mobile-CRISPRi (MCi) plasmid pJQ47 (pJMP2632, Addgene 134646, which has the sgRNA expression under control of the pTrc promoter and *Homo sapiens* codon-optimized *S. pyogenes* dCas9 expression under control of the Anderson BBa_J23117 weak constitutive promoter (17).

For individual gene construction, the plasmid vector was purified using the Purelink HiPure Plasmid Midiprep kit (Invitrogen K210005) and digested with BsaI-HFv2 (NEB R3733). Two 24-nucleotide oligonucleotides (“top” and “bottom”) encoding an sgRNA were designed with appropriate overhangs for Golden Gate Assembly. Oligos (2 µM each) were separately treated with T4 polynucleotide kinase and then annealed in 1X CutSmart buffer (NEB) at 95°C for 5 min followed by cooling to room temperature. The annealed insert (2 µl of a 1:40 dilution) was ligated into 50 ng BsaI-digested vector using T4 DNA ligase (NEB M0202). Plasmids were transformed into electrocompetent pir+ *E. coli* strain BW25141 and purified using the GeneJet Plasmid Miniprep kit (Thermo K0503). Following sequence confirmation, plasmids were transformed into electrocompetent pir+ *E. coli* mating strain WM6026.

Pooled CRISPRi plasmid libraries were cloned as described previously with minor modifications (46). A pooled sgRNA library covering genes in Supplementary Table S4 was ordered as single-stranded DNA oligonucleotides (Twist Bioscience) and generated by PCR amplification using Q5 DNA polymerase with the following components per 100 µl reaction: 20 µl Q5 buffer, 3 µl GC enhancer, 2 µl 10mM each dNTPs, 5 µl each 10 µM primers oJMP852 and oJMP853, 2 µl 10 nM oligonucleotide library, and 1 µl Q5 DNA polymerase (NEB M0491). Thermocycling conditions were: 98°C for 30s; 15 cycles of 98°C for 15s, 56°C for 15s, 72°C for 15s; followed by final extension at 72°C for 10 min. PCR products were digested with BsaI-HFv2 and ligated into BsaI-digested MCi plasmid(18). The ligation was purified by spot dialysis on nitrocellulose filter against 0.1 mM Tris, pH 8 buffer for 20 min. The library was transformed into *E. coli* BW25141, achieving approximately 90,000 colonies (∼30x coverage) across multiple plates. Colonies were pooled, plasmid DNA extracted, and transformed into WM6026 mating strain, yielding approximately 300,000 colonies (∼100x coverage). Library stocks were normalized to OD600 = 8 in LB with DAP and 15% glycerol for storage at -80°C.

### Transfer to P. aeruginosa PA14

The Mobile-CRISPRi system was transferred to PA14 through tri-parental mating as described for individual guides (18) and for pooled library construction (46) with minor modifications. Donor strains (*E. coli* WM6026 containing either the CRISPRi plasmid or Tn7 transposase) were grown in LB with ampicillin and DAP, while the PA14 recipient was grown in LB without supplements. Cultures were normalized to OD600 ∼3 and mixed in equal proportions (100 µl of each strain for individual constructs; 1.2 ml of each strain for library construction). Mixed cells were spotted onto cellulose filters (13 mm for individual constructs, 25 mm for library; MF-Millipore HAWG01300 or HAWG02500) and incubated at 37°C for 5 hours. Cells were recovered and plated on selective media (LB with gentamicin, no DAP). For library construction, approximately 240,000 colonies (∼80x coverage) were collected. Library stocks were normalized to OD600 = 10 in LB with 15% glycerol for storage at -80°C.

### Mouse infection with pooled Mobile-CRISPRi library

Starting with the resuspension of the glycerol stock in PBS, the inocula were prepared with two serial ten-fold dilutions. The more concentrated of the two inocula was diluted and spread on PIA and PIA + 30 μg/mL gentamicin plates for CFU enumeration. The remaining contents of the glycerol tube were centrifuged, and the pellet was frozen for gDNA extraction.

Pathogen-free male C57BL/6J mice at 8 weeks of age were purchased from Jackson Laboratories. Animal experiments were conducted in accordance with the approval of the Institutional Animal Care and Use Committee (IACUC) at UCSF. A total of 10 mice were anesthetized with isofluorane prior to intratracheal instillation with 50 μl of the *Pseudomonas* knockdown library per an established protocol (22). Animal weights and rectal temperatures were measured at every 6 hours to monitor the course of the infection.

Mice were sacrificed 24 hours post-infection. Lungs were collected in 3 ml of sterile PBS and homogenized by grinding the lung tissue against a cell strainer with the back of a syringe plunger under sterile conditions. 100 μL of lung homogenates were directly plated on 10 PIA + 30 μg/mL gentamicin plates. Then the homogenates were diluted to various degrees in LB media and the same dilution was plated on both PIA and PIA + 30 μg/mL gentamicin plates for CFU enumeration. The plates were incubated for 48 hours prior to harvest. 3-6 mL of LB was used to scrape colonies off PIA + 30 ug/mL gentamicin plates with an L-shaped spreader, and the 10 plates were combined to generate each mouse sample. These cell suspensions were centrifuged and stored at - 80 °C prior to gDNA extraction.

### Mouse infection with single strains

In separate experiments, 5 mL ON cultures of *ispD-P1*, *pgsA-P1*, *mrfp-P1*, and *WT* single-strain knockdown mutants in LB +/- 30 μg/mL gentamicin were grown from glycerol stocks at 37 °C with shaking at 225 rpm. After 16 hours, cultures were diluted 1:100 in 3 mL LB +/- 30 μg/mL gentamicin and were incubated with shaking until OD_600nm_ measured 0.64 (approximately 3 hours). 1 mL of the sub-culture was washed and resuspended in 1 mL PBS. The suspensions were diluted according to predetermined calculations based on OD_600nm_ measurements to yield a target inoculum of 2E6 CFU/animal. Each group contained 6 to 8 C57BL/6 mice at 8 to 12 weeks of age obtained from Jackson Laboratory, and the prior mouse infection protocol was followed. Survival was measured over 96 hours with temperature and weight measured every 6 hours.

### Amplicon library preparation & analysis

Qiagen DNeasy Blood & Tissue kit was used to extract gDNA from samples, and NEBNext Ultra II Q5® Master Mix was used for amplicon library preparation. Custom-made TruSeq primers extend the amplicon to incorporate the i5 and i7 ends, which are recognized by DualSeq primers procured from the Chan-Zuckerburg Biohub. The DualSeq primers are indexed to indicate sample identity and were demultiplexed after NGS. To determine number of reads needed from NGS, the number of unique barcodes was multiplied by a factor of 1,500 (3,112 * 1,500 = ∼5,000,000 reads) for robust detection of strain depletion.

### Growth Curves

3 mL LB + 30 μg/mL gentamicin cultures were inoculated with each PA14 strain and incubated at 37 °C with shaking at 225 rpm for 16 hours. Cultures were diluted 1:100 into fresh LB media and 200 μL of the respective cultures was added to each well in a 96-well plate. This plate was covered with an optically clear seal, and a needle was used to poke holes in each of the wells. OD_600nm_ were monitored during incubation in a microplate reader (Synergy H1; BioTek Instruments, VT) with continuous, fast, double orbital shaking. Samples were blanked with a well containing LB media. Results are representative of three technical replicates and at least two biological replicates.

### Guide RNA Sequence Abundance Analysis

Sequencing data were filtered to include samples with >5 million reads and >85% mapping rate. Analysis was performed using edgeR (v4.2.0) with a quasi-likelihood negative binomial model framework, comparing three conditions: initial inoculum, in vitro growth, and in vivo growth (high-inoculum samples only, due to increased noise in low-inoculum conditions). Control guides served as reference for baseline corrections. For genes with multiple guides, significance estimates were combined using Stouffer’s method and effect sizes were summarized using median log-fold changes. Statistical significance was assessed using quasi-likelihood F-tests with FDR correction for multiple comparisons. Data analysis and visualization were performed using R (packages: edgeR v4.2.0, data.table v1.15.4, ggplot2 v3.5.1).

### Population Bottleneck Estimations

Population bottleneck sizes were estimated using frequency-based calculations as described in Abel, et al. (22). Guide frequencies were compared between the initial inoculum and each subsequent condition to estimate the effective population size (*N*b) maintained through each transition. Analysis was performed separately for control and knockdown guides. The average *N*b values were calculated across replicates for each condition, with error bars representing standard error of the mean. This analysis provides quantitative estimates of the population complexity preserved throughout different stages of the experiment.

### Gene-Set Enrichment Analysis

Functional enrichment analysis was performed using predicted genome interactions from STRING-DB (v12.0.2) using the P. aeruginosa UCBPP-PA14 proteome (accessible at https://version-12-0.string-db.org/organism/STRG0A01FJP). Guide-level differential abundance results were analyzed using camera (limma v3.60.0) to perform competitive gene set testing, with control guides serving as the background set. Gene sets were derived from STRING-DB functional annotations including Gene Ontology terms, protein domains, and pathway annotations. Results were filtered based on false discovery rate (FDR) with enrichment direction and magnitude reported for each significant term.

## Supplementary Information

**Supplementary Figure 1: Construction of a *Pseudomonas* essential gene knockdown library**

The Mobile-CRISPRi library consists of 3,112 sgRNAs, including 1,000 non-targeting control sgRNAs and 4 targeting sgRNAs for each of 528 PA14 genes previously identified as essential in LB or infection-relevant media. sgRNAs were cloned into Mobile-CRISPRi vectors containing a constitutive P1 promoter driving dCas9 expression. The library was transformed into an *E. coli* donor strain and chromosomally integrated into PA14 through triparental mating to generate the pooled CRISPRi strain library.

**Supplementary Figure 2. sgRNA distribution and guide recovery during Mobile-CRISPRi library construction and sequencing**

**(A)** Density plots showing the distribution of sgRNA counts (Counts per Million) for non-targeting (black) and gene-targeting (red) guides in the *E. coli* mating strain and the PA14 transconjugant.

**(B)** Guide recovery per gene across experimental conditions including *E. coli* donor strain, PA14 transconjugant, LB cultures, and murine infection inocula (high and low dose). Table shows the number of genes with 0, 1, 2, 3, 4, and >0 guides detected in each condition.

**Supplementary Figure 3. Competitive gene-set enrichment analysis.**

**(A-C)** Competitive gene-set enrichment analysis using CAMERA for **(A)** Folate biosynthesis (KW-0289, 2/2 genes present), **(B)** Isoprene biosynthesis (CL:924, 5/5 genes present), and **(C)** Arginine biosynthesis (CL:2388, 5/5 genes present) pathways. Log2 fold change for guides targeting genes in these pathways is shown for in vitro vs. initial and in vivo vs. in vitro comparisons, with FDR values noted below each condition. Boxplots represent the distribution of log2 fold changes, and red lines indicate the significance-weighted median for each condition, when the gene-set enrichment is significant (FDR ≤ 0.05). The size of each point reflects guide-level significance (false discovery rate, FDR).

**Table S1. sgRNA sequences and targeted genes in knockdown library design**

Complete list of sgRNA sequences and their corresponding target genes used in the CRISPRi library construction.

**Table S2. Data from *in vitro* and *in vivo* screens normalized to inoculum**

Log_2_ fold change and false discovery rates for all detected hypomorphs and the statistical comparison between conditions.

**Table S3. Biological Pathways**

Guide-level CAMERA analysis results and statistics for all biological pathways tested in the CRISPRi screen.

**Table S4. Strains, Plasmids and Oligonucleotides**

List of strains, plasmids, and oligonucleotides used in this study.

